# Synergistic Combination of Bioengineered MicroRNA and Chemotherapy Across High-Risk Neuroblastoma Subtypes

**DOI:** 10.64898/2026.09.30.755443

**Authors:** Yunyoung Lee, Albatool AlKhazal, Kathleen E. Doyle, Yu-Rim Ahn, Ana Maria Sandoval Castallanos, Mei-Juan Tu, Ai-Ming Yu, Jinhwan Kim, Erin G. Brown

## Abstract

Despite intensive treatment, high-risk neuroblastoma (HRNB) remains a leading cause of cancer-related mortality in children. Current treatment paradigms include multimodal systemic treatments and multi-agent chemotherapy, which is limited by substantial acute and long-term toxicities. Platinum-based chemotherapy is a cornerstone of this treatment but is fraught with side effects and stands to benefit from dose reduction strategies. MicroRNA (miR)-based therapeutics represent an attractive strategy to simultaneously moderate multiple oncogenic pathways, providing a potential avenue to enhance the efficacy of current treatment regimens. However, the development of miR therapeutics has largely relied on chemically synthesized ones, which are limited by issues of inconsistency and poor in vivo stability. Biologically produced, tRNA/pre-miR containing bioengineered miRs (bioRNAs) address these limitations. We investigated the therapeutic potential of bioRNA encoding miR-124 and miR-34a across molecularly distinct HRNB models and evaluated their ability to enhance cisplatin efficacy. Both bioRNAs significantly suppressed neuroblastoma cell viability in vitro, with proteomic analyses demonstrating preferential induction of neuronal differentiation by miR-124 and apoptotic signaling by miR-34a. Combining bioRNA with cisplatin produced robust synergy across multiple HRNB lines, enhancing apoptosis while simultaneously promoting differentiation-associated phenotypes. In vivo, both bioRNAs suppressed tumor growth as monotherapies, while combination with low-dose cisplatin produced the strongest responses across multiple xenograft models with different HRNB subtypes. Notably, low-dose cisplatin combined with bioRNA therapy frequently achieved tumor control comparable to, or greater than, that observed with high-dose cisplatin monotherapy while maintaining favorable tolerability. These findings establish bioRNAs as potent cisplatin sensitizers and support bioRNA-based combination therapy as a strategy for improving HRNB treatment while reducing chemotherapy-associated toxicity in children.

## INTRODUCTION

Neuroblastoma arises from the aberrant development of the sympathetic nervous system and represents the most common extracranial solid tumor in young children.^1–5^ Although favorable-risk cases can often be cured, high-risk neuroblastoma (HRNB) remains the leading cause of pediatric cancer mortality in young children.^1–4^ HRNB is characterized by aggressive local disease, early metastasis, and pronounced molecular heterogeneity, driven by complex genetic and epigenetic alterations.^3,4,6^ These rapid tumor adaptations result in the development of therapeutic resistance and relapse observed in patients, which remains a significant predictor of mortality.^3,6,10^ Furthermore, one of the most clinically important molecular features is MYCN amplification, which defines a particularly aggressive subset of HRNB, although substantial therapeutic resistance and disease progression also occur in MYCN-non-amplified tumors. These biologically distinct subtypes can differ in differentiation state, apoptotic susceptibility, and response to therapy, posing a major challenge for treatment directed against individual molecular targets.^3,6,10^

Current treatment regimens for HRNB involve intensive multimodal systemic approaches, including high-dose chemotherapy, immunotherapy, and stem cell transplantation in addition to local control measures such as surgery and radiation. ^4,7,10^ However, these strategies are associated with significant systemic toxicity, a critical concern in pediatric patients whose developing organ systems are particularly vulnerable to long-term damage.^4,8^ Treatment-related toxicities, such as neurocognitive impairment, end-organ toxicity, ototoxicity, infertility, and secondary malignancies, pose lifelong risks. Therefore, there is a pressing need for innovative therapeutic strategies that enhance efficacy of current treatments while minimizing treatment-associated toxicity.^4,8^

MicroRNA (miRNA)-based therapeutics offer a promising strategy to address these challenges.^9–12^ miRNAs are endogenous regulators of gene expression that orchestrate posttranscriptional gene regulation (PTGR) across multiple signaling pathways.^10,13^ In neuroblastoma, dysregulation of tumor-suppressive miRNAs contributes to oncogenic signaling, impaired differentiation, and resistance to apoptosis.^9,18,19^ Restoration of such miRNAs has been shown to suppress tumor growth and reprogram tumor cell states.^20–22^ In particular, miR-124 and miR-34a, known to regulate differentiation, apoptosis, and tumor suppression pathways.^20–22^ Importantly, the ability of miRNAs to modulate gene networks, rather than individual targets, provides a conceptual advantage in addressing the intrinsic heterogeneity of HRNB.^10,13^ We therefore sought to determine whether restoration of miR-124 and miR-34a could produce antitumor effects across biologically distinct HRNB models, including MYCN-amplified and MYCN-non-amplified. In addition to their direct regulatory effects, miRNA-based therapeutics have the potential to synergize with conventional chemotherapy. By sensitizing tumor cells to cytotoxic stress, miRNA restoration can enhance therapeutic response and enable dose reduction of chemotherapeutic agents.^15,16^ This strategy is particularly appealing in pediatric oncology, where minimizing long-term toxicity is as important as achieving tumor eradication.^15^

Nonetheless, the clinical translation of miRNA therapeutics has been hampered by challenges linked to conventional chemically synthesized RNA molecules, including instability, inefficient delivery, off-target effects, and immunogenicity.^14,17^ Recently, biologically derived RNA platforms have emerged as an alternative.^23,24^ Among these, our recent work demonstrated an RNA bioengineering platform enabling the production of "true biologic RNA" (bioRNA) molecules with enhanced stability and functional compatibility in living systems.^23–25^ Given the molecular heterogeneity of HRNB, we hypothesized that bioRNA-mediated delivery of miR-124 and miR-34a could provide both direct antitumor activity across distinct HRNB subtypes while also enhancing sensitivity to chemotherapy, both in vitro and in vivo. To test this concept across diverse disease backgrounds, we evaluated bioRNA therapeutics in Be(2)-C, a MYCN-amplified HRNB model; SK-N-SH, a MYCN-non-amplified neuroblastoma model; and NB1643, a patient-derived MYCN-amplified HRNB model. Across all of these models, bioRNA treatment showed direct antitumor effects and enhanced the response to chemotherapy. Strikingly, combination treatment with bioRNA and chemotherapy facilitates effective tumor control at half-dose chemotherapy, suggesting a strategy to mitigate systemic toxicity while preserving therapeutic efficacy. These results establish a framework for integrating bioRNA-based miRNA therapeutics with existing treatment regimens and highlight their potential to overcome tumor heterogeneity and drug resistance in HRNB.

## RESULTS

### Bioengineered miR-124 and miR-34a suppress neuroblastoma growth across molecularly distinct HRNB models

To evaluate the therapeutic potential of bioRNA encoding tumor-suppressive miRs, miR-124 and miR-34a, in HRNB, we first assessed cell viability following treatment with either bioRNA (bioRNA-miR124 or bioRNA-miR34a) across a panel of biologically distinct HRNB cell lines, including SK-N-SH, Be(2)-C, and NB1643. These models represent genetically and phenotypically diverse HRNB subtypes and therefore provide an opportunity to assess the robustness of bioRNA-mediated tumor suppression across heterogeneous disease backgrounds.

Treatment with either bioRNA-miR124 or bioRNA-miR34a significantly reduced cell viability relative to untreated, vehicle, and scrambled RNA controls **(Fig. 1A)**. Although the magnitude of response varied among cell lines, both bioRNAs consistently suppressed NB cell growth. Notably, SK-N-SH and NB1643 cells exhibited particularly strong responses, with viability reduced to approximately 30–40% of control levels following both bioRNA treatment, whereas Be(2)-C cells displayed a more moderate but still significant reduction in viability. Similar trends were observed across a range of cisplatin concentrations, indicating that the growth-suppressive effects of both bioRNAs were maintained under conditions of chemotherapeutic stress **(Supplementary Fig. 1)**.

**Figure 1.**
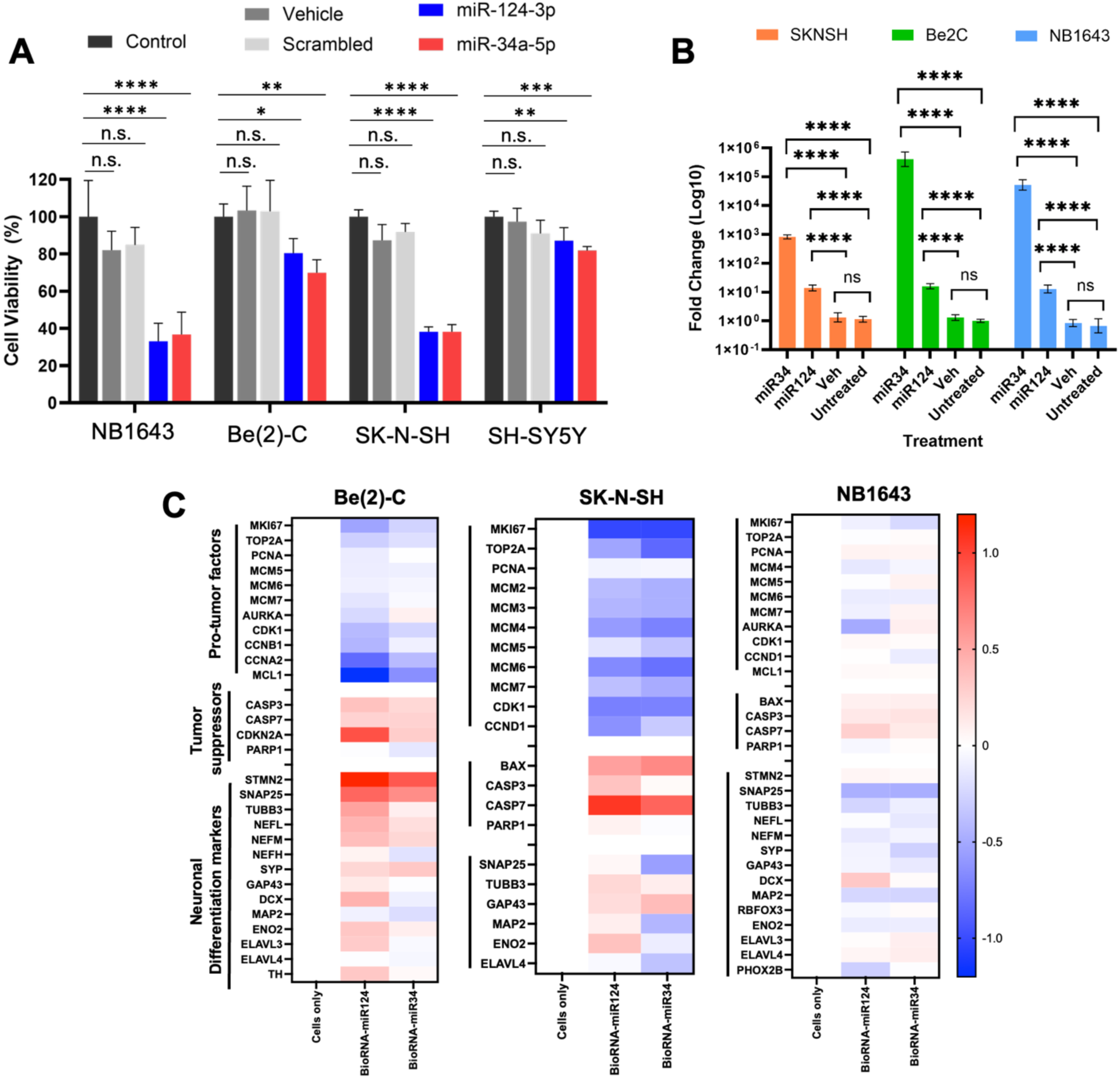
Bioengineered miR-124 and miR-34a suppress neuroblastoma growth through distinct differentiation- and apoptosis-associated pathways. **(A)** Cell viability following treatment with vehicle, scrambled RNA control, bioengineered miR-124, or bioengineered miR-34a at a fixed concentration. **(B)** RT-qPCR quantification of intracellular miR-124 and miR-34a levels following treatment in neuroblastoma cell lines. Data are presented as mean ± SEM. Statistical significance was determined using one-way ANOVA with appropriate multiple-comparison correction. **(C)** Quantitative analysis showing differential regulation of proliferation-associated proteins, apoptosis-associated proteins, and neuronal differentiation markers following bioRNA treatment.

To verify successful intracellular delivery and processing of the bioRNA construct, mature miR expression levels were quantified by RT-qPCR. Robust intracellular accumulation of mature miR-124 and miR-34a was observed in all three cell lines, with expression levels increasing by several orders of magnitude relative to untreated controls **(Fig. 1B)**. These results demonstrate efficient cellular uptake and processing of the bioRNA constructs into mature miRNAs. While RT-qPCR does not directly assess biological activity, the marked increase in mature miRNA levels establishes a basis for subsequent functional and mechanistic analyses.

To investigate molecular pathways associated with bioRNA-mediated growth suppression, we performed proteomic profiling following bioRNA-miR124 and bioRNA-miR34 treatment, respectively. Based on our analysis of approximate 5,100 proteins, both bioRNAs produced antitumor effects, while they induced distinct biological programs **(Fig. 1C)**. Using a threshold of log_2_FC > 0.5, bioRNA-miR124 altered 1,492 proteins (32.2%) in SK-N-SH, 1,032 proteins (20.8%) in Be(2)C, and 385 proteins (7.9%) in NB1643. BioRNA-miR34a altered 1,588 (34.3%), 573 (11.3%), and 540 (11.2%) proteins in the respective cell lines. Across multiple cell lines, bioRNA-miR124 treatment preferentially increased proteins associated with neuronal differentiation, including TUBB3, SNAP25, MAP2, DCX, and other markers of neuronal maturation. These changes were most evident in SK-N-SH (moderate) and Be(2)-C cells (strong), suggesting that miR-124 promotes partial differentiation of NB cells toward a less proliferative cellular state. In contrast, bioRNA-miR34a treatment produced a stronger apoptotic signature. Proteins associated with programmed cell death, including BAX, CASP3, CASP7, and PARP1, were consistently elevated following miR-34a exposure. Concurrently, several proliferation-associated proteins, including MKI67, TOP2A, PCNA, CDK1, and CCND1, were reduced, supporting a shift from active proliferation toward apoptotic cell fate commitment. Interestingly, the magnitude of proteomic remodeling differed among cell lines. SK-N-SH and Be(2)-C cells displayed pronounced alterations in both differentiation- and apoptosis-related pathways, whereas NB1643 cells exhibited comparatively weaker proteomic changes despite demonstrating substantial reductions in cell viability. These findings suggest that multiple mechanisms may contribute to bioRNA-mediated tumor suppression and further underscore the biological heterogeneity of HRNB.

Collectively, these data demonstrate that bioRNA-miR124 and bioRNA-miR34a exert potent antitumor activity across genetically diverse neuroblastoma models while engaging distinct differentiation and apoptotic programs.

### BioRNA treatment cooperates with cisplatin to enhance antitumor activity

Because cisplatin remains a central component of current NB therapy, we next investigated whether restoration of tumor-suppressive miRNA signaling via bioRNA treatment could enhance cisplatin responsiveness. HRNB cells were treated with combinations of bioRNAs and cisplatin, and drug interactions were analyzed using Highest Single Agent (HSA)-based synergy modeling. Across all three HRNB cell lines, combined treatment produced greater growth inhibition than either agent alone **(Fig. 2A, B)**. Synergy mapping revealed positive interaction scores throughout broad regions of the dose-response landscape rather than being restricted to isolated concentration pairs **(Fig. 2C, D)**. Particularly strong synergy was observed in Be(2)-C and SK-N-SH cells, where large portions of the response surface exhibited positive HSA scores, indicating cooperative interactions between bioRNA treatment and cisplatin. Although NB1643 cells displayed comparatively lower synergy scores, positive interactions remained detectable across multiple concentration combinations, suggesting that bioRNA-mediated sensitization is conserved across distinct HRNB subtypes.

**Figure 2.**
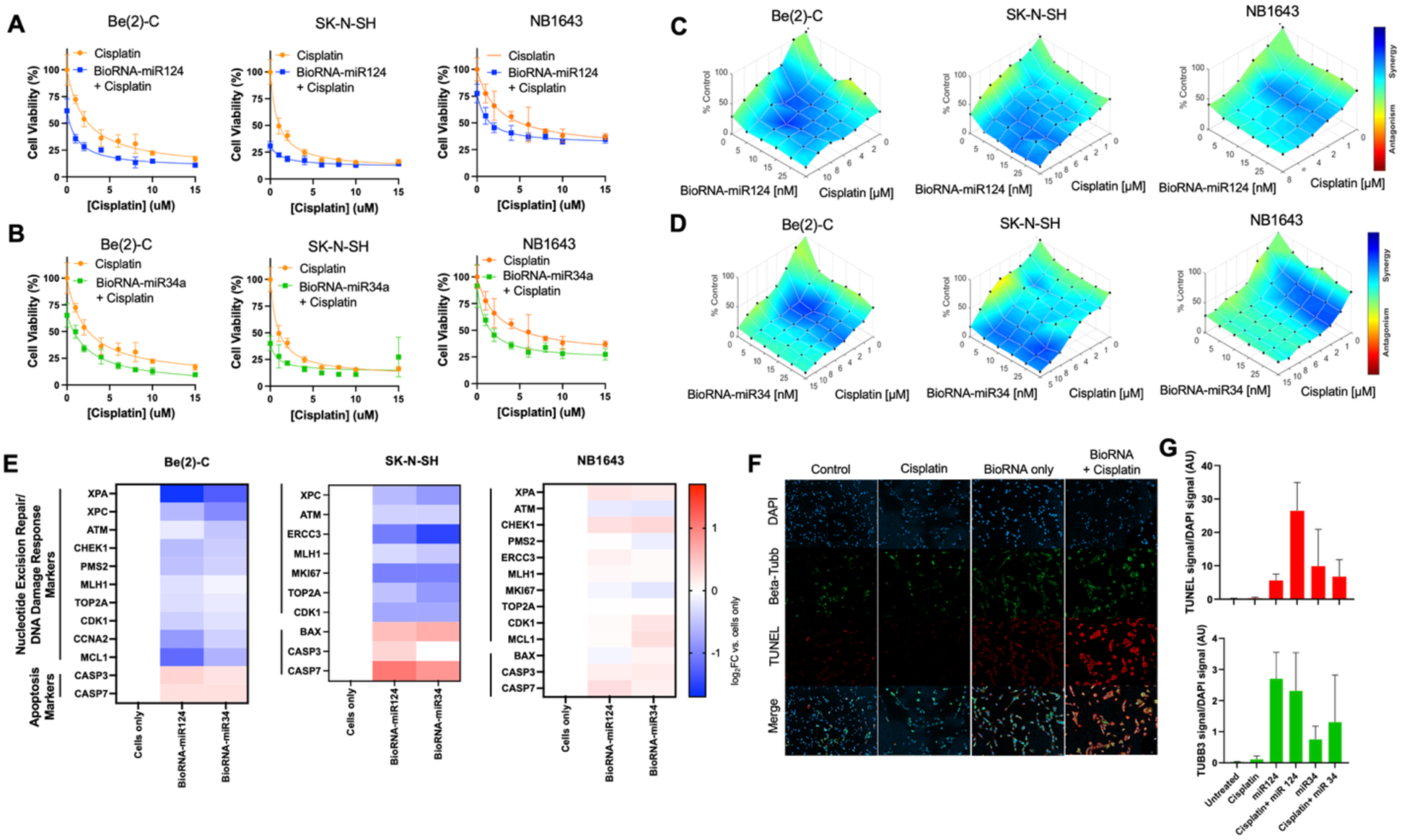
BioRNA treatment synergizes with cisplatin through modulation of DNA damage repair and cell fate pathways. **(A-B)** Cell viability following treatment with cisplatin in the presence or absence of bioengineered miR-124 **(A)** or miR-34a **(B)** in Be(2)-C, SK-N-SH, and NB1643 cells. **(C-D)** Highest Single Agent (HSA)-based synergy analysis demonstrating cooperative interactions between bioRNA-miR124 **(C)** or bioRNA-miR-34a **(D)** with cisplatin across multiple concentration combinations in Be(2)-C, SK-N-SH, and NB1643 cells. **(E)** Quantitative proteomic analysis of proteins associated with nucleotide excision repair (NER), homologous recombination-mediated DNA damage repair (HR-DSB), cell-cycle regulation, and apoptosis following bioRNA treatment. **(F)** Representative immunofluorescence images of SK-N-SH cells stained for βIII-tubulin (green), TUNEL (red), and nuclei (blue). **(G)** Quantification of TUNEL (top) and βIII-tubulin (bottom) signals normalized to DAPI staining. Data are presented as mean ± SD.

To explore mechanisms underlying this synergistic response, we analyzed key proteins involved in DNA damage signaling and repair pathways. The results showed that several key DNA repair proteins following bioRNA treatment were highly suppressed **(Fig. 2E)**. Specifically, components of nucleotide excision repair (NER), including XPA and XPC, were reduced, particularly in Be(2)-C and NB1643 cells. Additional decreases were observed in ATM, CHEK1, PMS2, MLH1, and ERCC3, proteins that participate in DNA damage recognition, checkpoint activation, and repair of cisplatin-induced lesions. In contrast, pro-apoptotic proteins including BAX, CASP3, and CASP7 were elevated. Because cisplatin exerts its cytotoxic activity primarily through induction of DNA crosslinks and genotoxic stress, suppression of adaptive DNA repair pathways would be expected to increase cellular susceptibility to chemotherapy-induced apoptosis. The observed molecular alterations therefore provide a possible mechanistic explanation for the synergistic interactions observed between bioRNA treatment and cisplatin.

To validate these findings at the cellular level, we performed immunofluorescence analyses using TUNEL staining and βIII-tubulin (TUBB3) staining as indicators of apoptosis and neuronal differentiation, respectively (**Fig. 2F, G and Supplementary Fig. 2**). In SK-N-SH cells, combination treatment induced the highest levels of TUNEL positivity, consistent with enhanced apoptotic cell death. In parallel, bioRNA-miR124-containing treatment groups exhibited increased βIII-tubulin expression relative to untreated controls, supporting activation of neuronal differentiation. Similar trends were observed in Be(2)-C cells, where bioRNA-miR124-treated groups generated the strongest differentiation signals while bioRNA-miR34a-treated groups produced pronounced apoptotic responses but not strong neuronal differentiation, in consistency with our proteomic analysis as shown in **Fig. 1B**.

Together, these findings indicate that bioRNA-miR124 and bioRNA-miR34a sensitize NB cells to cisplatin through complementary biological mechanisms involving differentiation induction, apoptotic activation, and attenuation of DNA repair pathways.

### BioRNA-mediated sensitization to cisplatin induces durable tumor regression in SK-N-SH xenografts without evidence of systemic toxicity

Next, to evaluate the therapeutic efficacy of bioRNA-miR124 and bioRNA-miR34a in vivo, we first examined whether either bioRNA could suppress tumor growth as a monotherapy in SK-N-SH NB xenograft as a representative MYCN-non-amplified in vivo model. We then asked whether bioRNA treatment could enhance the efficacy of cisplatin and thereby support chemotherapy dose de-escalation, which is an important consideration in the treatment of HRNB because of the substantial acute and long-term toxicities associated with intensive chemotherapy. Mice were treated with bioRNA-miR124 or bioRNA-miR34a alone, low-dose cisplatin (10 mg/kg), high-dose cisplatin (20 mg/kg), or a combination of bioRNA with low-dose cisplatin (10 mg/kg).

Tumors in untreated animals exhibited rapid and progressive growth throughout the study period, reaching greater than 30-fold expansion relative to baseline **(Fig. 3A)**. Both miR-124 and miR-34a monotherapies significantly delayed tumor progression compared with controls, confirming that bioRNAs retain intrinsic antitumor activity in vivo. Cisplatin treatment also suppressed tumor growth in a dose-dependent manner, with 20 mg/kg producing greater inhibition than 10 mg/kg. Addition of either bioRNA to low-dose cisplatin produced substantially greater tumor suppression than either monotherapy and consistently outperformed low-dose and high-dose cisplatin alone. Notably, the bioRNA-miR124/cisplatin combination induced complete tumor regression in 4 of 6 animals (67%), while the bioRNA-miR34a/cisplatin combination achieved complete regression in 3 of 6 animals (50%). These high rates of complete regression were particularly notable given that the combination groups received only the low dose of cisplatin.

**Figure 3.**
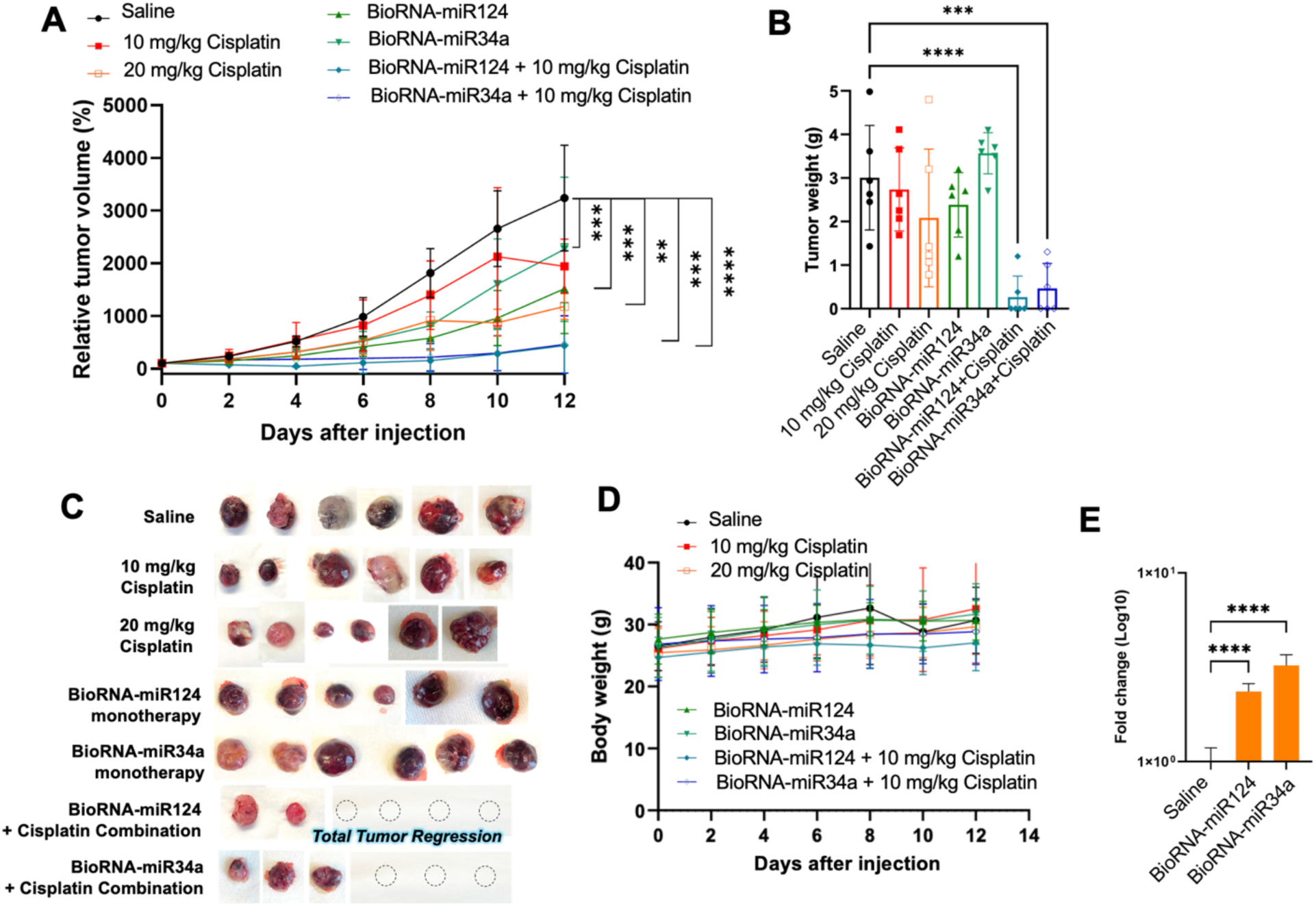
BioRNA-cisplatin combination therapy induces durable tumor regression and maintains systemic tolerability in MYCN-nonamplified SK-N-SH xenografts. **(A)** Relative tumor growth curves following treatment with bioRNAs, cisplatin, or combination therapy. **(B)** Endpoint tumor weights. **(C)** Representative gross images of harvested tumors. Dashed circles indicate complete tumor regression. **(D)** Body weight measurements throughout treatment. **(E)** RT-qPCR analysis of intratumoral miR-124 and miR-34a levels at study termination. Data are presented as mean ± SD.

Consistent with longitudinal tumor measurements, endpoint tumor weights were significantly reduced in both combination groups **(Fig. 3B)**. Gross examination of harvested tumors further demonstrated marked reductions in tumor burden, and complete tumor regression was observed in several animals receiving the bioRNA-miR124/cisplatin combination **(Fig. 3C).** Analysis of individual tumor growth trajectories demonstrated that the therapeutic benefit was broadly distributed across animals and was not driven by isolated responders **(Supplementary Fig. 3)**.

To confirm successful intratumoral delivery of bioRNAs, RT-qPCR analysis was performed on excised tumors. Both miR-124 and miR-34a accumulated robustly within tumor tissues, demonstrating efficient in vivo delivery and persistence throughout the treatment period **(Fig. 3E).** Histopathological examination provided additional evidence supporting treatment efficacy. H&E staining revealed increased regions of tumor degeneration and necrosis following treatment, particularly in combination-treated tumors **(Supplementary Fig. 4).** Extensive necrotic areas were especially evident in the miR-34a plus cisplatin group, indicating profound disruption of tumor viability and tissue architecture.

Importantly, enhanced antitumor activity was not associated with overt systemic toxicity. Body weight remained stable throughout the study across all treatment groups, including combination-treated animals **(Fig. 3D)**. To further evaluate treatment tolerability, serum chemistry analyses were performed at study termination. No substantial alterations were observed in biomarkers associated with renal function (BUN), hepatic injury (ALT and ALP), metabolic status (glucose), or serum protein levels (albumin and globulin) among treatment groups compared with controls **(Supplementary Fig. 5)**. Collectively, these findings indicate that bioRNA-mediated enhancement of cisplatin efficacy can be achieved without detectable systemic toxicity under the conditions tested.

### BioRNA-cisplatin combination therapy demonstrates broad efficacy in MYCN-amplified Be(2)-C neuroblastoma

To determine whether the therapeutic benefit of bioRNA-based treatment extended across biologically distinct HRNB models, we next evaluated treatment efficacy in MYCN-amplified Be(2)-C xenografts. Similar to the SK-N-SH model, untreated tumors displayed rapid growth kinetics throughout the study period **(Fig. 4A)**. However, the response to bioRNA monotherapy differed from that observed in SK-N-SH tumors. Neither bioRNA-miR124 nor bioRNA-miR34a alone exhibited a clear reduction in tumor growth, as shown in longitudinal tumor volume, endpoint tumor weight, and gross tumor images **(Fig. 4A, B, C)**. This differential response highlights the variable sensitivity of molecularly distinct HRNB models to miR restoration alone, consistent with our in vitro observation that Be(2)-C cells were relatively less responsive to bioRNA monotherapy **(Fig. 1A)**. Cisplatin monotherapy also did not show significant tumor regression, possibly due to the highly aggressive nature of MYCN-amplified Be(2)-C model even at the high-dose treatment group. Despite the limited effect of bioRNA monotherapy and cisplatin monotherapy, addition of either bioRNA to low-dose cisplatin markedly enhanced tumor progression relative to untreated controls. Although complete regressions were not observed as shown in SK-N-SH tumors, both bioRNA-cisplatin combinations significantly outperformed corresponding monotherapies.

**Figure 4.**
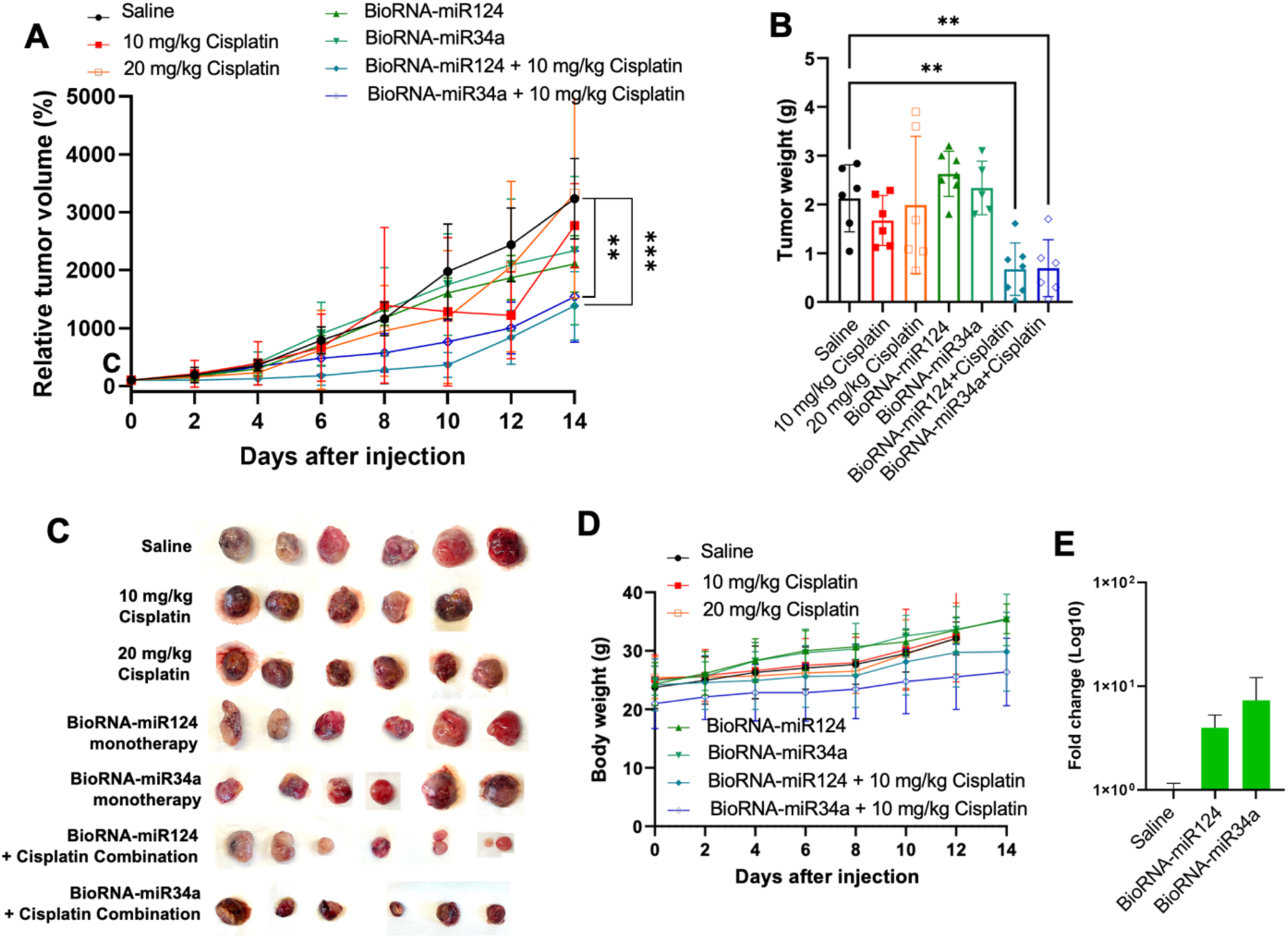
BioRNA-mediated sensitization enhances cisplatin efficacy in MYCN-amplified Be(2)-C neuroblastoma xenografts. **(A)** Relative tumor growth curves during treatment. **(B)** Endpoint tumor weights. **(C)** Representative images of excised tumors. **(D)** Body weight measurements throughout the study. **(E)** RT-qPCR quantification of intratumoral miR-124 and miR-34a expression levels following treatment. Data are presented as mean ± SD.

Consistent with longitudinal tumor measurement, endpoint tumor weights were significantly reduced in animals receiving combination therapy and represented the lowest values among all treatment groups **(Fig. 4B).** Gross pathological examination confirmed these findings, revealing substantially smaller tumors in combination-treated animals compared with control or monotherapy groups **(Fig. 4C)**. Individual tumor growth curves demonstrated that therapeutic responses were consistently observed across animals and were not limited to a subset of exceptional responders **(Supplementary Fig. 6)**.

RT-qPCR analysis confirmed successful intratumoral delivery of both bioRNAs and production of corresponding miRs following repeated administration **(Fig. 4E)**. Furthermore, body weights remained stable throughout treatment and did not differ substantially among groups, suggesting that combination therapy was well tolerated despite enhanced antitumor efficacy **(Fig. 4D)**. Lastly, H&E staining demonstrated increased tumor degeneration and necrotic regions in combination-treated tumors, with the bioRNAs plus cisplatin groups exhibiting extensive tissue destruction **(Supplementary Fig. 7)**. Together, these findings demonstrate that bioRNA-mediated enhancement of cisplatin efficacy at lower dose and their synergy is strong in a representative MYCN-amplified and highly aggressive NB subtype, Be(2)-C, which is not responding well to monotherapy.

### Potent antitumor responses are maintained in NB1643 xenografts following bioRNA-cisplatin combination treatment

Lastly, we evaluated therapeutic efficacy in NB1643 patient-derived HRNB cell line xenografts. In NB1643 xenograft model, which is also MYCN-amplified, the therapeutic response toward each treatment was slightly different than other HRNB models, Untreated tumors expanded rapidly throughout the study period, and low dose cisplatin slightly slow down the tumor growth yet not statically significant than control. In contrast, bioRNA monotherapy and high-dose cisplatin monotherapy exhibited significant delay in tumor progression **(Fig. 5A)**. Combination treatment again generated the strongest therapeutic response. Both bioRNA-miR124 plus cisplatin and bioRNA-miR34a plus cisplatin produced sustained suppression of tumor growth and maintained tumor volumes substantially below those observed in control animals and other monotherapy groups throughout the study. Notably, the bioRNA-miR34a and cisplatin combination displayed particularly strong activity and nearly abolished tumor progression in several animals, with one mouse showed total regression.

**Figure 5.**
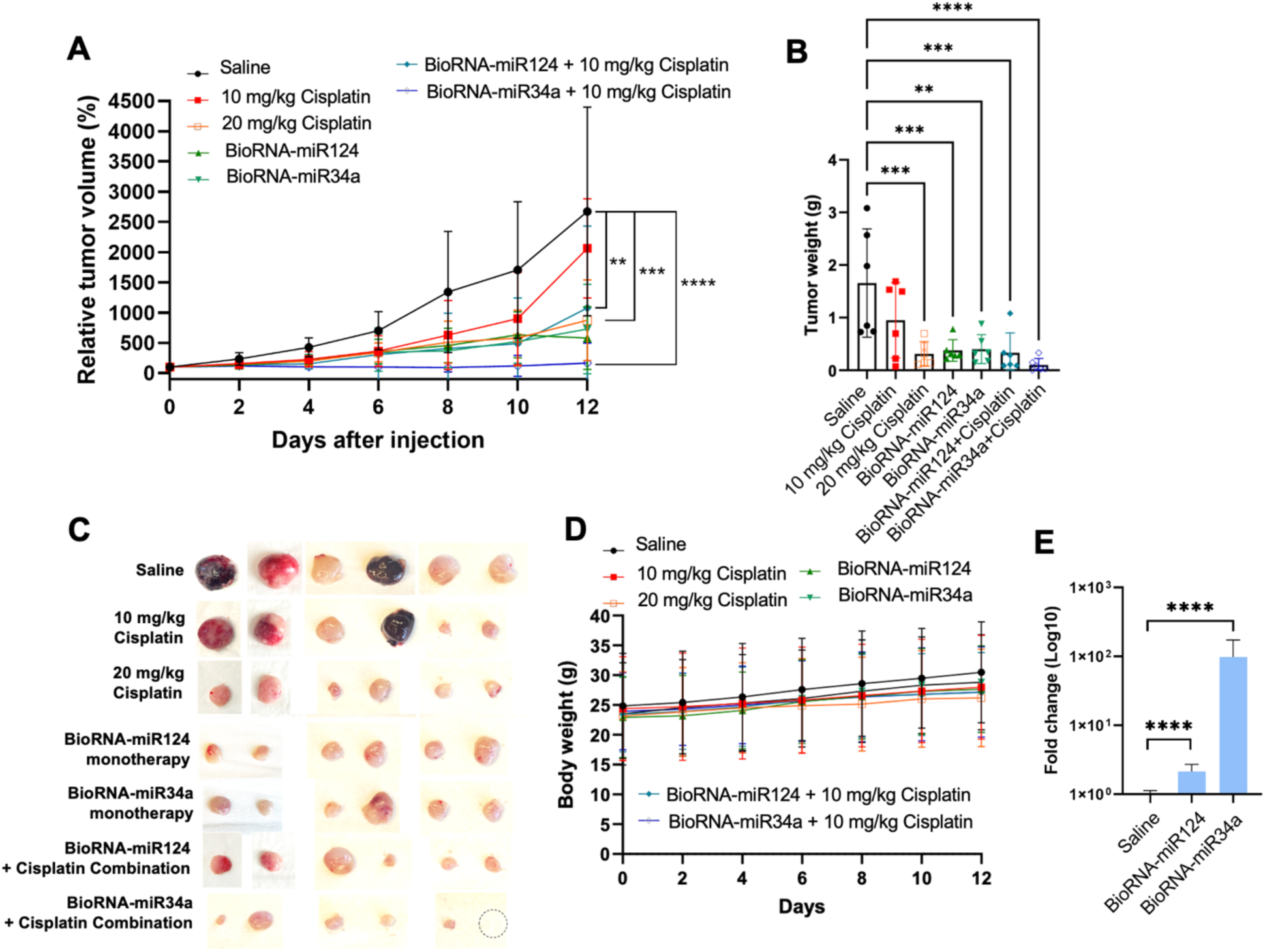
Potent antitumor activity of bioRNA-cisplatin combination therapy in patient-derived NB1643 neuroblastoma xenografts. **(A)** Relative tumor growth curves during treatment. **(B)** Endpoint tumor weights. **(C)** Representative images of harvested tumors. Dashed circles indicate complete tumor regression. **(D)** Body weight measurements throughout treatment. **(E)** RT-qPCR analysis of miR-124 and miR-34a accumulation in tumor tissues at study endpoint. Data are presented as mean ± SD.

These effects were confirmed by endpoint tumor weight measurements, which demonstrated significant reductions in tumor burden in all combination-treatment groups relative to controls and monotherapies **(Fig. 5B)**. Gross examination of harvested tumors further highlighted the magnitude of treatment response, revealing dramatically smaller residual tumors following combination treatment **(Fig. 5C)**. Analysis of individual tumor growth trajectories revealed highly consistent treatment responses across animals, supporting the reproducibility of the observed antitumor effects **(Supplementary Fig. 8)**. RT-qPCR analysis confirmed efficient delivery and retention of both miRNAs within tumor tissues, with particularly robust accumulation of miR-34a observed at study termination **(Fig. 5E)**. Importantly, body weights remained stable throughout treatment and did not indicate treatment-associated toxicity despite profound tumor regression **(Fig. 5D)**. These findings demonstrate that bioRNA-mediated chemotherapy sensitization remains highly effective in NB1643 tumors and further support the broad applicability of this therapeutic approach across diverse HRNB subtypes.

## DISCUSSION

HRNB remains one of the most challenging pediatric malignancies because of its extensive biological heterogeneity, high relapse rates, and poor outcomes despite aggressive multimodal therapy.^1–8^ Although platinum-based chemotherapy remains a key component of multimodal treatment, cumulative cisplatin exposure is associated with substantial short- and long-term toxicities that can significantly impact quality of life in survivors.^4,8^ Consequently, therapeutic strategies capable of enhancing antitumor efficacy while reducing chemotherapy burden are of considerable clinical interest. In the present study, we demonstrate that bioRNA-miR124 and bioRNA-miR34a possess intrinsic antitumor activity and, importantly, significantly enhance cisplatin responsiveness across multiple neuroblastoma models. Most notably, low-dose cisplatin combined with bioRNA treatment consistently achieved tumor control comparable to or greater than that observed with high-dose cisplatin monotherapy, supporting the potential of bioRNA-mediated chemotherapy sensitization as a strategy to improve therapeutic index and lower the chemotherapy dose for HRNB treatment in children.

Our findings extend our previous studies demonstrating the therapeutic potential of bioengineered RNA agents. Unlike conventional synthetic oligonucleotide approaches, bioRNAs are generated through biological processes that more closely resemble endogenous RNA biosynthesis and processing.^23–25^ Our present study demonstrates that this platform can successfully restore tumor-suppressive miRNA function in HRNB and, importantly, can be leveraged to enhance the efficacy of existing chemotherapeutic agents. From a translational perspective, the ability to integrate bioRNA therapy with current standard-of-care regimens may facilitate improved clinical chemotherapeutic response while minimizing disruption to established treatment paradigms.

A key finding of this study is that both bioRNAs suppress HRNB growth through distinct yet complementary biological mechanisms. Proteomic analyses revealed that bioRNA-miR124 preferentially induced neuronal differentiation-associated proteins, including markers linked to neuronal maturation and reduced proliferative potential.^27–29^ These observations are consistent with previous reports identifying miR-124 as a central regulator of neuronal differentiation and neural lineage commitment.^22^ Because NB originates from incompletely differentiated neural crest-derived precursor cells, restoration of differentiation programs represents an attractive therapeutic strategy. The increased βIII-tubulin expression observed following bioRNA-miR124 treatment further supports the notion that it suppresses tumor growth, at least in part, by promoting a more differentiated and less aggressive cellular phenotype. In contrast, bioRNA-miR34a induced a stronger apoptotic response characterized by increased expression of pro-apoptotic proteins and activation of cell death pathways.^30–31^ These findings are consistent with previous studies identifying miR-34a as a potent tumor suppressor frequently downregulated in neuroblastoma and capable of inhibiting tumor progression through modulation of multiple oncogenic signaling networks.^20,21^ Interestingly, despite their mechanistic differences, both bioRNAs produced substantial antitumor activity in vitro and in vivo. This observation suggests that restoration of distinct tumor-suppressive programs may ultimately converge on a common therapeutic outcome, namely suppression of tumor growth and increased vulnerability to cytotoxic stress.

One of the most significant findings of this study is the robust synergy observed between bioRNA treatment and cisplatin. Combination analyses demonstrated positive interactions across broad concentration ranges in all three HRNB cell lines tested, indicating that the observed cooperation is unlikely to be restricted to isolated drug concentrations. Instead, bioRNA-mediated sensitization appears to represent a generalizable biological phenomenon. Consistent with this observation, combination treatment produced superior antitumor efficacy in every xenograft model examined, including SK-N-SH, Be(2)-C, and NB1643 tumors. Mechanistically, our protein-level biomarker analyses suggest that attenuation of DNA repair pathways may contribute substantially to this synergistic response. Multiple proteins involved in nucleotide excision repair (NER), DNA damage recognition, mismatch repair, and checkpoint signaling were reduced following miRNA treatment. Because NER is one of the principal pathways responsible for removing cisplatin-induced DNA adducts, suppression of these repair mechanisms would be expected to increase tumor cell sensitivity to platinum-mediated DNA damage.^26,27^ Together, these findings support a model in which restoration of tumor-suppressive miRNAs lowers the threshold for irreversible cell death following chemotherapy-induced DNA damage.

An additional finding that warrants consideration is the differential relationship between molecular response and therapeutic efficacy among neuroblastoma models. SK-N-SH and Be(2)-C cells exhibited pronounced proteomic remodeling following miRNA treatment, whereas NB1643 cells demonstrated comparatively weaker changes in several pathway-level analyses. Despite this difference, NB1643 tumors remained highly responsive to bioRNA-cisplatin combination therapy as well as bioRNA monotherapy in vivo. This apparent disconnect suggests that extensive proteomic remodeling may not be strictly required for therapeutic response and raises the possibility that a limited number of critical regulatory nodes are sufficient to drive cell apoptosis and chemotherapy sensitization.^33^

The broad efficacy observed across multiple HRNB models is particularly encouraging given the extensive intertumoral heterogeneity characteristic of HRNB. NB encompasses diverse genetic and epigenetic states, and many targeted therapies demonstrate efficacy only within restricted molecular subsets.^3,6^ In contrast, both bioRNAs maintained activity across MYCN-amplified and non-amplified models despite substantial differences in baseline sensitivity and molecular responses.^32,34^ This broad activity likely reflects one of the fundamental advantages of bioRNA/miRNA-based therapeutics: the ability to simultaneously regulate multiple interconnected pathways rather than targeting a single oncogenic driver.^9–13^ Such network-level regulation may be particularly valuable in heterogeneous malignancies where pathway redundancy frequently limits the durability of highly specific targeted therapies.

Several limitations should be acknowledged. First, the current studies were performed using subcutaneous xenograft models and therefore do not fully recapitulate the complex tumor microenvironment or metastatic behavior of human NB. Second, bioRNAs were administered locally, whereas future clinical translation will likely require systemic delivery strategies capable of achieving efficient tumor targeting and biodistribution. Third, although our data identify DNA repair pathways as candidate mediators of therapeutic synergy, additional mechanistic studies will be required to define the direct downstream targets responsible for chemotherapy sensitization. Future investigations should evaluate orthotopic and metastatic disease models, advanced delivery systems, and potential interactions between bioRNA therapy and emerging immunotherapeutic approaches.

## CONCLUSION

In conclusion, we demonstrate that bioRNA-miR124 and bioRNA-miR34a function as potent antitumor agents and chemotherapy sensitizers in HRNB. By promoting differentiation, activating apoptosis, and attenuating DNA repair pathways, bioRNA treatment enhances cisplatin responsiveness across multiple HRNB subtypes. Most importantly, bioRNA-mediated sensitization enables effective tumor control using reduced cisplatin doses, frequently achieving outcomes comparable to or exceeding those obtained with high-dose cisplatin monotherapy. These findings support further development of bioRNA-based combination strategies aimed at improving therapeutic efficacy while minimizing treatment-associated toxicity in pediatric NB.

## MATERIALS AND METHODS

### Materials

Eagle’s Minimum Essential Medium (EMEM, **30-2003**) and Iscove’s Modified Dulbecco’s Medium (IMDM, 30-2005) were purchased from American Type Culture Collection (ATCC). Ham’s F 12 Nutrient Mix (F-12, 11-765-054), Dulbecco’s Phosphate Buffered Saline (DPBS, SH3002803), Fetal Bovine Serum (FBS, FB12999102), **Pen**icillin-Streptomycin Solution, 100X, 10000 U/mL **Pen**icillin, 10000 ug/mL Streptomycin (SV30010), Trypsin (MT2505CI), Opti-MEM (31-985-062), **Lipofectamine** 3000 Transfection Reagent (L3000008), Insulin-Transferrin-Selenium (ITS, MT25800CR) were purchased form Fisher Scientific

### Neuroblastoma Cell Culture

Three human HRNB cell lines were used: NB1643 (Childhood Cancer Repository, Lubbock, TX, USA), Be(2)-C, and SK-N-SH (American Type Culture Collection [ATCC], Manassas, VA, USA). SK-N-SH cells were cultured in EMEM; ATCC supplemented with 10% FBS and 1% penicillin–streptomycin. Be(2)-C cells were maintained in a 1:1 mixture of EMEM and Ham’s F-12 Nutrient Mix, supplemented with 10% FBS and 1% penicillin–streptomycin. NB1643 cells were cultured in IMDM supplemented with 20% FBS, 1% penicillin–streptomycin, and 1% ITS. All cell lines were maintained at 37°C in a humidified incubator with 5% CO₂.

### Bioengineering of BioRNA agents

All bioengineered RNA (bioRNA) agents, including miR-124-3p and miR-34a-5p, as well as a control RNA consisting of human glycyl tRNA fused to a Sephadex aptamer (GSA), were generated and obtained from the laboratory of Ai-Ming Yu in the Department of Biochemistry and Molecular Medicine at the University of California, Davis School of Medicine (Sacramento, CA, USA). Reagents were procured through the Stratocore PPMS software via the Molecular Pharmacology Shared Resource (MPSR) at UC Davis.

The expression and purification of bioengineered miR-124 and miR-34 have been previously described by Ai-Ming Yu and colleagues [1, 2, 3, 4].

### Cell viability assay

Cell viability was assessed using the 3-(4,5-dimethylthiazol-2-yl)-2,5-diphenyltetrazolium bromide (MTT) assay (Sigma-Aldrich, M2128). High-risk neuroblastoma (HRNB) cells were seeded at a density of 1× 10^4^ cells per well in 96-well plates and cultured overnight. The following day, cells were treated with the indicated conditions. For bioRNA treatments, cells were transfected with 25 nM bioengineered RNAs using Lipofectamine 3000 according to the manufacturer’s instructions. For chemotherapy treatment, cells were exposed to cisplatin. Cells were treated with either bioRNA alone, cisplatin alone, or combination treatment of bioRNA and cisplatin. Cells were exposed to the appropriate treatment for 72 hours. Following treatment, media was removed and cells were incubated with MTT solution (5 mg/mL) for 3–4 hours at 37°C. Formazan crystals were dissolved in DMSO, and absorbance was measured at 570 nm using a SpectraMax i3 microplate reader (Molecular Devices). Cell viability values were normalized to untreated control groups.

Drug combination effects were evaluated using Combenefit software. Synergy was quantified using the Highest Single Agent (HSA) model, in which the observed effect of the drug combination is compared to the maximal effect of each single agent alone at the corresponding concentrations. Positive HSA scores indicate synergistic interactions, whereas negative values indicate antagonism.

### Proteomics Analysis

High-risk neuroblastoma (HRNB) cells were seeded in 60 mm dishes at a density of 8 × 10⁵ cells per dish for SK-N-SH and NB1643 cells, and 4 × 10⁵ cells per dish for Be(2)-C cells. Cells were allowed to adhere overnight and subsequently transfected with 25 nM bioRNA using Lipofectamine3000. At 48 hours post-transfection, cells were harvested and lysed for proteomic analysis.

Cell lysis was performed using a buffer consisting of 70% diethyl pyrocarbonate (DEPC)-treated water, 25% (w/v) sodium dodecyl sulfate (SDS; prepared from a 20% stock solution), and 5% ammonium bicarbonate (AmBic). Lysates were collected and submitted to the UC Davis Proteomics Core Facility for protein digestion and liquid chromatography–tandem mass spectrometry (LC–MS/MS) analysis.

### TUNEL staining and immunocytochemistry

B(e)2-C and SK-N-SH cells were seeded onto glass coverslips in 12-well plates at a density of 1 × 10⁵ cells per well and incubated for 24 hours. Cells were then treated with the appropriate bioRNA or cisplatin and incubated for 6 hours. Following treatment, media was removed and cells were washed once with DPBS, then fixed in neutral buffered formalin for 15 minutes at room temperature.

Apoptosis was assessed using the Click-iT™ Plus TUNEL Assay Kit for *In Situ* Apoptosis Detection (Fisher Scientific, C10618) according to the manufacturer’s protocol. Following the TUNEL reaction, immunofluorescence staining for βIII-tubulin was performed using an anti-βIII tubulin primary antibody (Abcam, ab18207) and a goat anti-rabbit IgG H&L secondary antibody (Abcam, ab150077), following the manufacturer’s instructions. Nuclei were counterstained with Hoechst 33342 (Fisher Scientific, H3570) for 15 minutes.

Coverslips were mounted onto microscope slides using ProLong™ antifade mounting medium (Fisher Scientific). Images were acquired using a Zeiss LSM 900 confocal microscope.

### RT-qPCR *in vitro*

HRNB cells were seeded at a density of 1 × 10⁵ cells per well in 12-well plates and transfected with miR-124-3p (25 nM), miR-34a-5p (25 nM), Lipofectamine 3000 alone, or left untreated as a control. Total RNA was extracted 48 hours post-transfection using the RNeasy Mini Kit (Qiagen, Germantown, MD, USA).

Complementary DNA (cDNA) was synthesized using the SuperScript III First-Strand Synthesis System (Invitrogen, Thermo Fisher Scientific, Cat. No. 18080051). Random hexamers were used for U6, while stem-loop primers were used for miRNA-specific reverse transcription: miR-124-3p (5′-GTCGTATCCAGTGCAGGGTCCGAGGTATTCGCACTGGATACGACGGCATT-3′) and miR-34a-5p (5′GTCGTATCCAGTGCAGGGTCCGAGGTATTCGCACTGGATACGACACAACC-3′). Quantitative real-time PCR was performed using a QuantStudio 5 Real-Time PCR System (Applied Biosystems, Waltham, MA, USA) with gene-specific primers. For miR-124-3p, the forward primer was 5′-GCGCTAAGGCACGCGGTG-3′ and the reverse primer was 5′-GTGCAGGGTCCGAGGT-3′. For miR-34a-5p, the forward primer was 5′-CGCGCTGGCAGTGTC TTAGCT-3′ and the reverse primer was 5′-GTGCAGGGTCCGAGGT-3′. U6 snRNA was used as an internal control with forward primer 5′-CTCGCTTCGGCAGCACA-3′ and reverse primer 5′-AACGCTTCACGAATTTGCGT-3′. Relative expression levels were calculated using the comparative Ct (2⁻ΔΔCt) method.

### In vivo validation

#### Experimental murine xenograft neuroblastoma mouse models and miR-124 and miR-34 therapy studies

All animal procedures were approved by the Institutional Animal Care and Use Committee (IACUC) at the University of California, Davis (UC Davis, USA). Five-to six-week-old female non-obese diabetic/severe combined immunodeficient (NOD/SCID) mice were obtained from The Jackson Laboratory (Bar Harbor, ME, USA).

Murine xenograft neuroblastoma models were established by subcutaneous injection of high-risk neuroblastoma (HRNB) cells. Briefly, each mouse was inoculated with 2.0 × 10⁶ cells suspended in 50 μL saline into the flank. Tumor growth was monitored every two days by measuring body weight and tumor size using calipers. Tumor volume was calculated using the formula:

tumor volume (mm³) = 0.5 × [length (mm) × width² (mm²)].

Treatment was initiated once tumors reached approximately 100 mm³ in volume. Mice were randomized into seven treatment groups (n = 6 mice per group) for each cell line: (1) untreated control, (2) miR-124 alone, (3) miR-34 alone, (4) high-dose cisplatin (20 mg/kg), (5) low-dose cisplatin (10 mg/kg), (6) miR-124 combined with low-dose cisplatin, and (7) miR-34 combined with low-dose cisplatin.

Bioengineered RNA treatments were formulated with in vivo-jetPEI (30 μg) and administered via intratumoral injection. Cisplatin was administered intravenously. Combination treatments were delivered simultaneously. All treatments were administered as a single dose, and tumor growth and body weight were monitored every 2–3 days for approximately 14 days.

At the study endpoint, mice were euthanized and tumors were excised, measured, and weighed. Tumor tissues were then flash-frozen in liquid nitrogen for downstream quantitative PCR (qPCR) analysis.

#### Estimation of miR levels after treatment and in vivo

RT–qPCR was performed to assess miR-124-3p and miR-34a-5p expression in neuroblastoma tumors following treatment with bioengineered RNAs, compared with untreated HRNB tumors. Tumors were flash-frozen immediately after harvest and stored at −80°C until RNA extraction. Total RNA was isolated using the Quick-RNA Miniprep Plus Kit (Zymo Research, Irvine, CA, USA). cDNA was synthesized using the SuperScript III First-Strand Synthesis System (Invitrogen, Thermo Fisher Scientific, Cat. No. 18080051) with miRNA-specific stem-loop primers or random hexamers, as appropriate. Quantitative PCR was performed on a QuantStudio 5 Real-Time PCR System using stem-loop–specific primers for miR-124-3p, miR-34a-5p, and U6 snRNA. Relative miRNA expression levels were determined using the comparative Ct (2^−ΔΔCt) method.

### Statistical analysis

All results are expressed as mean ± standard deviation (SD). Statistical comparisons were performed using one- or two-way analysis of variance (ANOVA) followed by Bonferroni post hoc tests or Student’s *t*-test (GraphPad Prism, version 10, San Diego, CA, USA). The probability value of *P* < 0.05 was considered as statistically significant.

## Notes

### Competing Interest Statement

The authors have declared no competing interest.

### Funding

- Hartwell Foundation: SP0A242412
- American Pediatric Surgical Association: A25-0973
- National Institute of Health: P30CA093373

## REFERENCES

1. Newman EA, Abdessalam S, Aldrink JH, Austin M, Heaton TE, Bruny J, et al; APSA Cancer Committee. Update on neuroblastoma. J Pediatr Surg. 2019 Mar;54(3):383–389. doi: 10.1016/j.jpedsurg.2018.09.004. Epub 2018 Sep 19. PMID: 30305231.

2. Davidoff AM. Neuroblastoma. Semin Pediatr Surg. 2012 Feb;21(1):2-14. doi: 10.1053/j.sempedsurg.2011.10.009. PMID: 22248965; PMCID: PMC3261589.

3. Louis CU, Shohet JM. Neuroblastoma: molecular pathogenesis and therapy. Annu Rev Med. 2015;66:49-63. doi: 10.1146/annurev-med-011514-023121. Epub 2014 Oct 27. PMID: 25386934; PMCID: PMC4418018.

4. Pinto NR, Applebaum MA, Volchenboum SL, Matthay KK, London WB, Ambros PF, et al. Advances in risk classification and treatment strategies for neuroblastoma. J Clin Oncol. 2015 Sep 20;33(27):3008–3017. doi: 10.1200/JCO.2014.59.4648. Epub 2015 Aug 24. PMID: 26304901; PMCID: PMC4567703.

5. Maris JM. Recent advances in neuroblastoma. N Engl J Med. 2010 Jun 10;362(23):2202–2211. doi: 10.1056/NEJMra0804577. PMID: 20558371.

6. Pugh TJ, Morozova O, Attiyeh EF, Asgharzadeh S, Wei JS, Auclair D, et al. The genetic landscape of high-risk neuroblastoma. Nat Genet. 2013 Mar;45(3):279–284. doi: 10.1038/ng.2529. PMID: 23334666.

7. Yu AL, Gilman AL, Ozkaynak MF, London WB, Kreissman SG, Chen HX, et al. Anti-GD2 antibody with GM-CSF, interleukin-2, and isotretinoin for neuroblastoma. N Engl J Med. 2010 Sep 30;363(14):1324–1334. doi: 10.1056/NEJMoa0911123. PMID: 20879881.

8. Oeffinger KC, Mertens AC, Sklar CA, Kawashima T, Hudson MM, Meadows AT, et al. Chronic health conditions in adult survivors of childhood cancer. N Engl J Med. 2006 Oct 12;355(15):1572–1582. doi: 10.1056/NEJMsa060185. PMID: 17035650.

9. Aravindan N, et al. MicroRNAs in neuroblastoma tumorigenesis, therapy resistance, and disease evolution. Cancer Drug Resist. 2019;2:1086–1105.

10. Esquela-Kerscher A, Slack FJ. Oncomirs—microRNAs with a role in cancer. Nat Rev Cancer. 2006 Apr;6(4):259–269. doi: 10.1038/nrc1840. PMID: 16557279.

11. Rupaimoole R, Slack FJ. MicroRNA therapeutics: towards a new era for the management of cancer and other diseases. Nat Rev Drug Discov. 2017 Mar;16(3):203–222. doi: 10.1038/nrd.2016.246. PMID: 28209991.

12. Chakraborty C, Sharma AR, Sharma G, Doss CGP, Lee SS. Therapeutic miRNA and siRNA: moving from bench to clinic as next generation medicine. Mol Ther Nucleic Acids. 2017 Jun 16;8:132–143. doi: 10.1016/j.omtn.2017.06.005. PMID: 28777947.

13. Bartel DP. MicroRNAs: target recognition and regulatory functions. Cell. 2009 Jan 23;136(2):215–233. doi: 10.1016/j.cell.2009.01.002. PMID: 19167326.

14. Winkle M, El-Daly SM, Fabbri M, Calin GA. Noncoding RNA therapeutics— challenges and potential solutions. Nat Rev Drug Discov. 2021 Aug;20(8):629–651. doi: 10.1038/s41573-021-00219-z. Epub 2021 Jun 18. PMID: 34145432; PMCID: PMC8212082.

15. Pottoo FH, Barkat MA, Harshita, Ansari MA, Javed MN, Sajid Jamal QM, et al. Nanotechnological based miRNA intervention in the therapeutic management of neuroblastoma. Semin Cancer Biol. 2021 Feb;69:100–108. doi: 10.1016/j.semcancer.2019.09.017. Epub 2019 Sep 25. PMID: 31562954.

16. Garofalo M, Croce CM. MicroRNAs: master regulators of drug resistance. Nat Rev Drug Discov. 2011 Nov;10(11):857–868. doi: 10.1038/nrd3700. PMID: 22015900.

17. Setten RL, Rossi JJ, Han SP. The current state and future directions of RNAi-based therapeutics. Nat Rev Drug Discov. 2019 Jun;18(6):421–446. doi: 10.1038/s41573-019-0017-4. PMID: 30723287.

18. Stallings RL. MicroRNA involvement in neuroblastoma. Clin Genet. 2009 Sep;76(3):313–321. doi: 10.1111/j.1399-0004.2009.01208.x. PMID: 19793049.

19. Maugeri M, Barbagallo D, Barbagallo C, Banelli B, Di Mauro S, Purrello F, et al. Altered expression of miRNAs and methylation of their promoters are correlated in neuroblastoma. Oncotarget. 2016 Dec 13;7(50):83330–83341. doi: 10.18632/oncotarget.13090. PMID: 27829219; PMCID: PMC5347773.

20. De Antonellis P, Carotenuto M, Vandenbussche J, De Vita G, Ferrucci V, Medaglia C, et al. Early targets of miR-34a in neuroblastoma. Mol Cell Proteomics. 2014 Aug;13(8):2114–2131. doi: 10.1074/mcp.M113.035808. Epub 2014 Jun 9. PMID: 24912852; PMCID: PMC4125741.

21. Welch C, Chen Y, Stallings RL. MicroRNA-34a functions as a potential tumor suppressor by inducing apoptosis in neuroblastoma cells. Oncogene. 2007 Jul 26;26(34):5017–5022. doi: 10.1038/sj.onc.1210293. Epub 2007 Feb 12. PMID: 17297439.

22. Silber J, Lim DA, Petritsch C, Persson AI, Maunakea AK, Yu M, et al. miR-124 regulates neuronal differentiation and tumorigenesis. Cancer Res. 2008 May 1;68(9):3739–3746. doi: 10.1158/0008-5472.CAN-07-6757. PMID: 18483254.

23. Li MM, Yu AM. Bioengineered RNA agents (BERAs) for RNAi-based therapy. Pharmacol Ther. 2015 Apr;148:74–86. doi: 10.1016/j.pharmthera.2014.11.002. PMID: 25448662.

24. Wang W, Yu AM. Bioengineered noncoding RNA agents: emerging therapeutics. Adv Drug Deliv Rev. 2018 Jan;132:3–15. doi: 10.1016/j.addr.2018.09.002. PMID: 30244027.

25. Yu AM, Pan YZ. Noncoding microRNAs: small RNAs play a big role in drug metabolism. Annu Rev Pharmacol Toxicol. 2012;52:315–335. doi: 10.1146/annurev-pharmtox-010611-134643. PMID: 21910627.

26. Matthay KK, Maris JM, Schleiermacher G, Nakagawara A, Mackall CL, Diller L, Weiss WA. Neuroblastoma. Nat Rev Dis Primers. 2016 Nov 10;2:16078. doi: 10.1038/nrdp.2016.78. PMID: 27830764.

27. Makeyev EV, Zhang J, Carrasco MA, Maniatis T. The MicroRNA miR-124 promotes neuronal differentiation by triggering brain-specific alternative pre-mRNA splicing. Mol Cell. 2007 Aug 3;27(3):435–48. doi: 10.1016/j.molcel.2007.07.015. PMID: 17679093; PMCID: PMC3139456.

28. He L, He X, Lim LP, de Stanchina E, Xuan Z, Liang Y, Xue W, Zender L, Magnus J, Ridzon D, Jackson AL, Linsley PS, Chen C, Lowe SW, Cleary MA, Hannon GJ. A microRNA component of the p53 tumour suppressor network. Nature. 2007 Jun 28;447(7148):1130–4. doi: 10.1038/nature05939. Epub 2007 Jun 6. PMID: 17554337; PMCID: PMC4590999.

29. Cole KA, Attiyeh EF, Mosse YP, Laquaglia MJ, Diskin SJ, Brodeur GM, Maris JM. A functional screen identifies miR-34a as a candidate neuroblastoma tumor suppressor gene. Mol Cancer Res. 2008 May;6(5):735–42. doi: 10.1158/1541-7786.MCR-07-2102. PMID: 18505919; PMCID: PMC3760152.

30. Agostini M, Knight RA. miR-34: from bench to bedside. Oncotarget. 2014 Feb 28;5(4):872–81. doi: 10.18632/oncotarget.1825. PMID: 24657911; PMCID: PMC4011589

31. Bansal M, Gupta A, Ding HF. MYCN and Metabolic Reprogramming in Neuroblastoma. Cancers (Basel). 2022 Aug 25;14(17):4113. doi: 10.3390/cancers14174113. PMID: 36077650; PMCID: PMC9455056.

32. Krystal J, Foster JH. Treatment of High-Risk Neuroblastoma. Children (Basel). 2023 Jul 28;10(8):1302. doi: 10.3390/children10081302. PMID: 37628301; PMCID: PMC10453838.

33. Griso AB, Acero-Riaguas L, Castelo B, Cebrián-Carretero JL, Sastre-Perona A. Mechanisms of Cisplatin Resistance in HPV Negative Head and Neck Squamous Cell Carcinomas. Cells. 2022 Feb 5;11(3):561. doi: 10.3390/cells11030561. PMID: 35159370; PMCID: PMC8834318.

34. Cole KA, Attiyeh EF, Mosse YP, Laquaglia MJ, Diskin SJ, Brodeur GM, Maris JM. A functional screen identifies miR-34a as a candidate neuroblastoma tumor suppressor gene. Mol Cancer Res. 2008 May;6(5):735–42. doi: 10.1158/1541-7786.MCR-07-2102. PMID: 18505919; PMCID: PMC3760152.

