## Supplementary Information for "Synergistic Combination of Bioengineered MicroRNA and Chemotherapy Across High-Risk Neuroblastoma Subtypes"

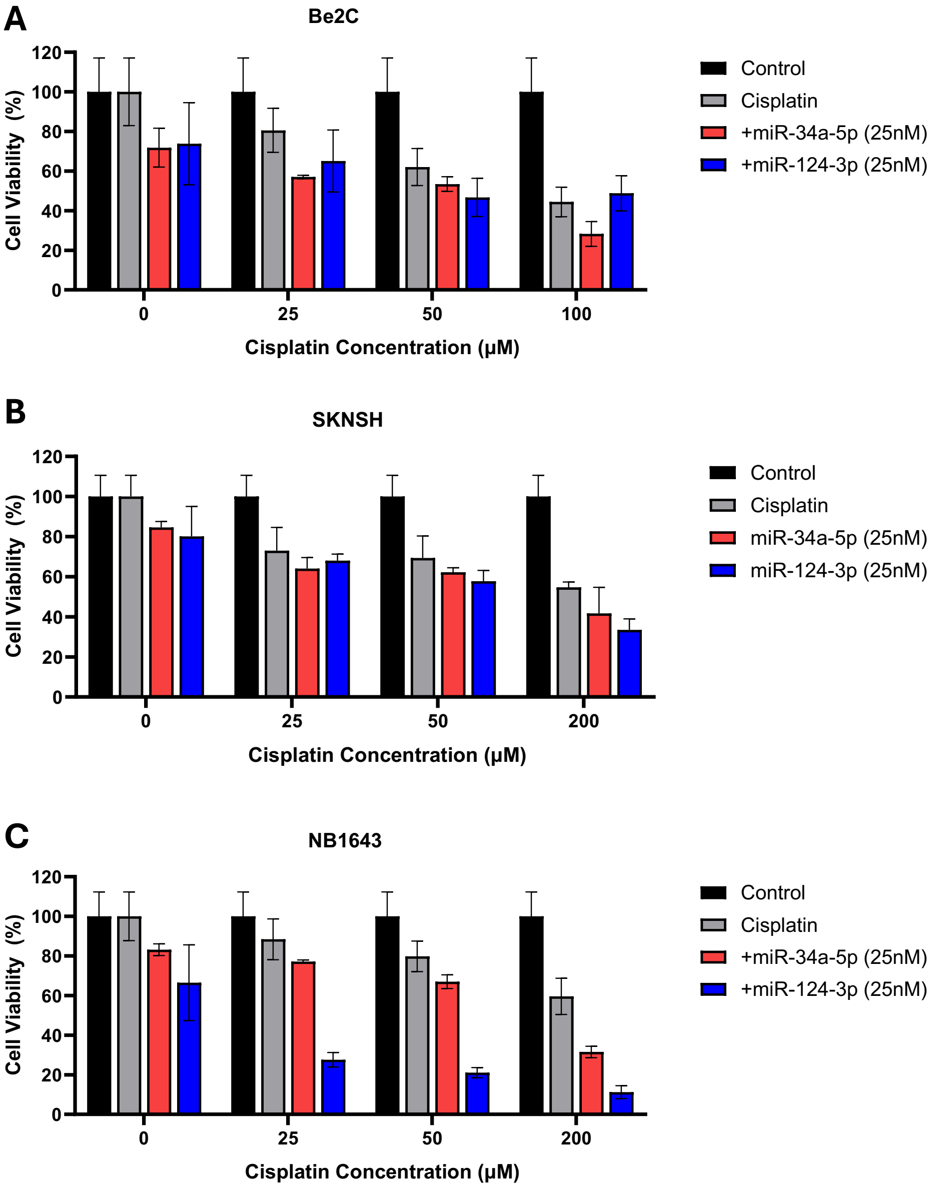


**Supplementary Figure 1. BioRNA treatment enhances cisplatin-mediated growth inhibition in neuroblastoma cell lines.**
(A–C) Cell viability of Be(2)-C, SK-N-SH, and NB1643 cells treated with increasing concentrations of cisplatin in the presence or absence of bioengineered miR-124 or miR-34a. Data are presented as mean ± SD.

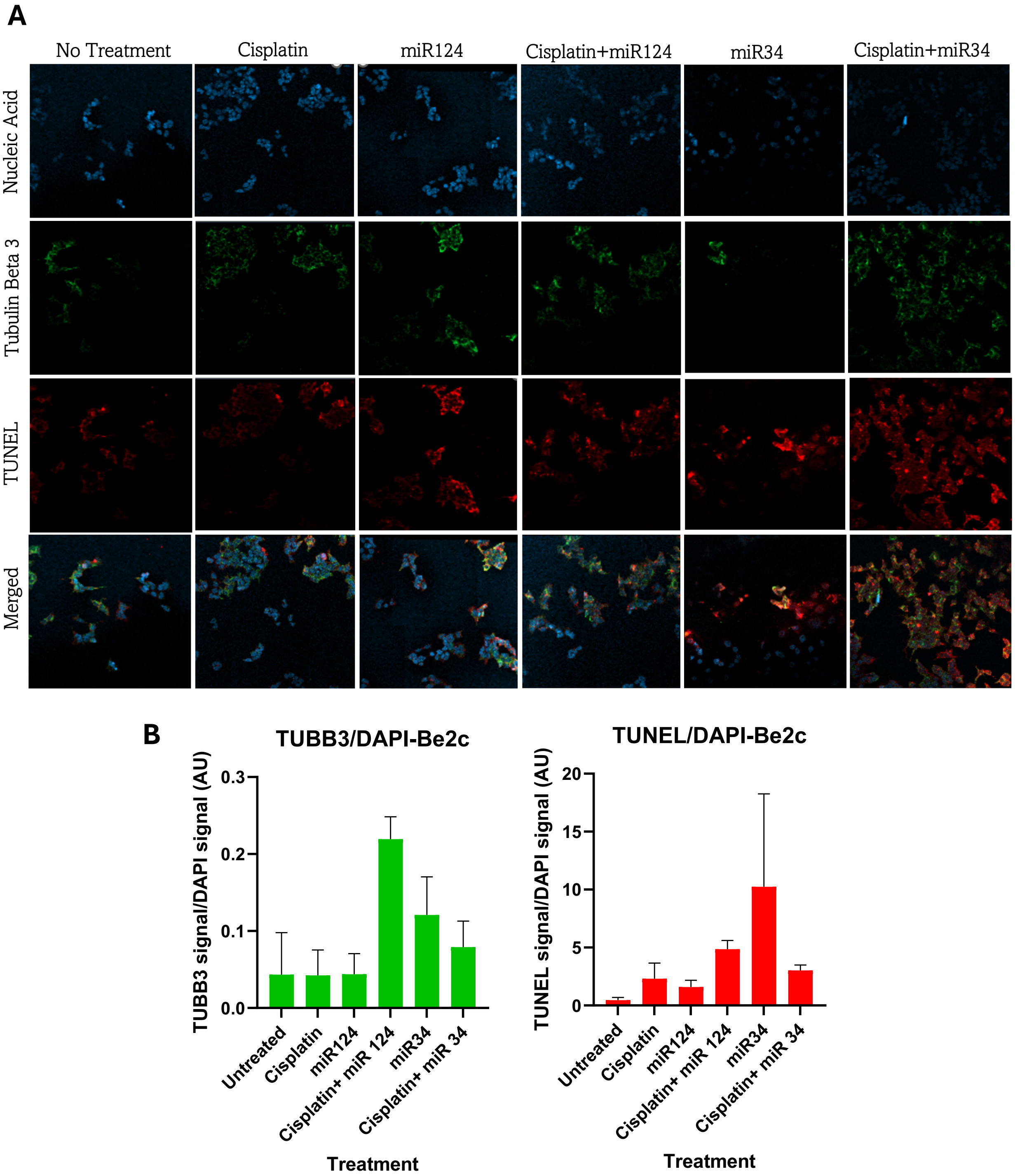


**Supplementary Figure 2. BioRNA treatment promotes apoptosis and neuronal differentiation in Be(2)-C cells.**

(A) Representative immunofluorescence images of Be(2)-C cells stained for βIII-tubulin (green), TUNEL (red), and nuclei (blue) following treatment with cisplatin, bioRNAs, or combination therapy. (B) Quantification of βIII-tubulin and TUNEL signals normalized to DAPI staining is shown below. Data are presented as mean ± SD.

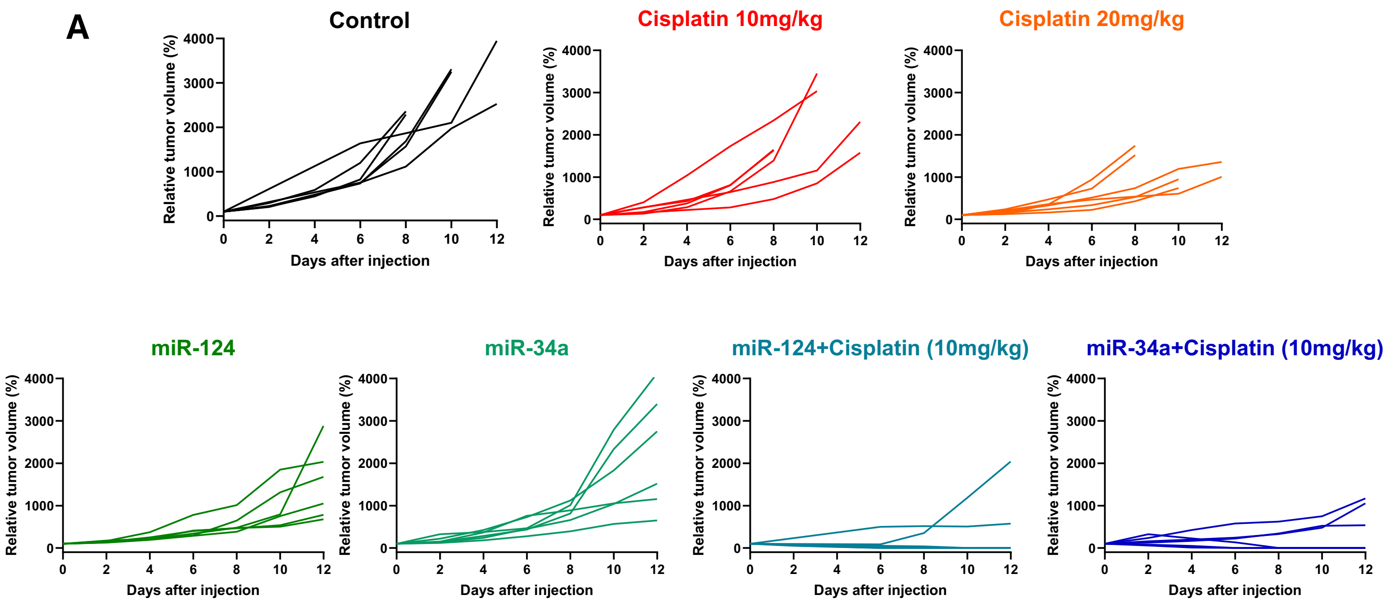


**Supplementary Figure 3. Individual tumor growth trajectories in SK-N-SH xenografts.**

(A) Spider plots showing relative tumor volume progression for individual animals receiving control, cisplatin, bioRNA monotherapy, or combination treatment.

A

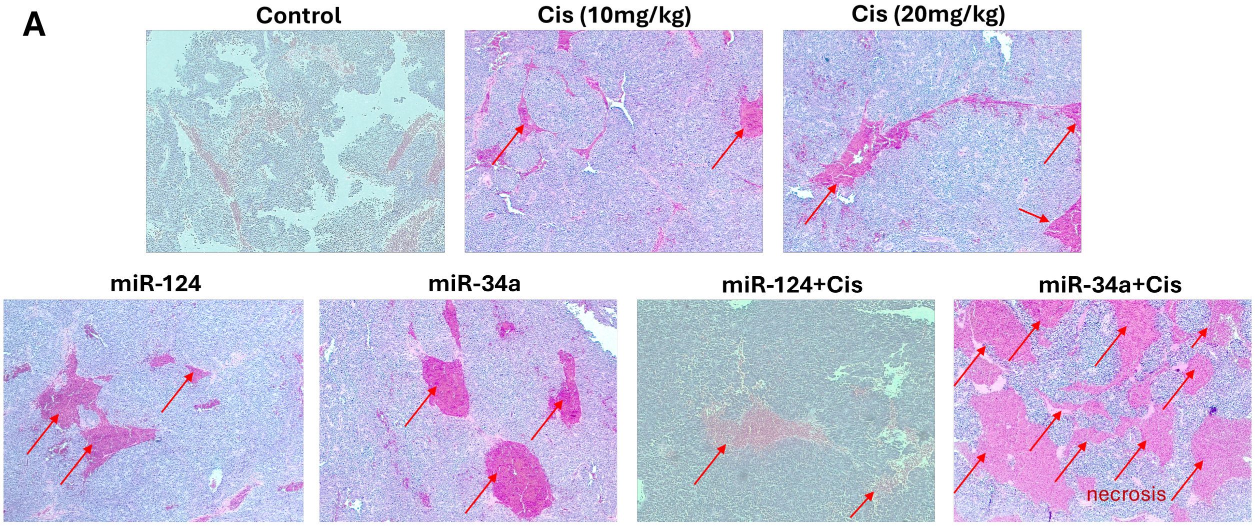


**Supplementary Figure 4. Histopathological evaluation of treatment responses in SK-N-SH xenografts.**
(A) Representative H&E-stained tumor sections collected at study endpoint. Arrows indicate regions of tissue degeneration and necrosis. Extensive necrotic areas were observed in tumors receiving bioRNA-cisplatin combination therapy.

**A**

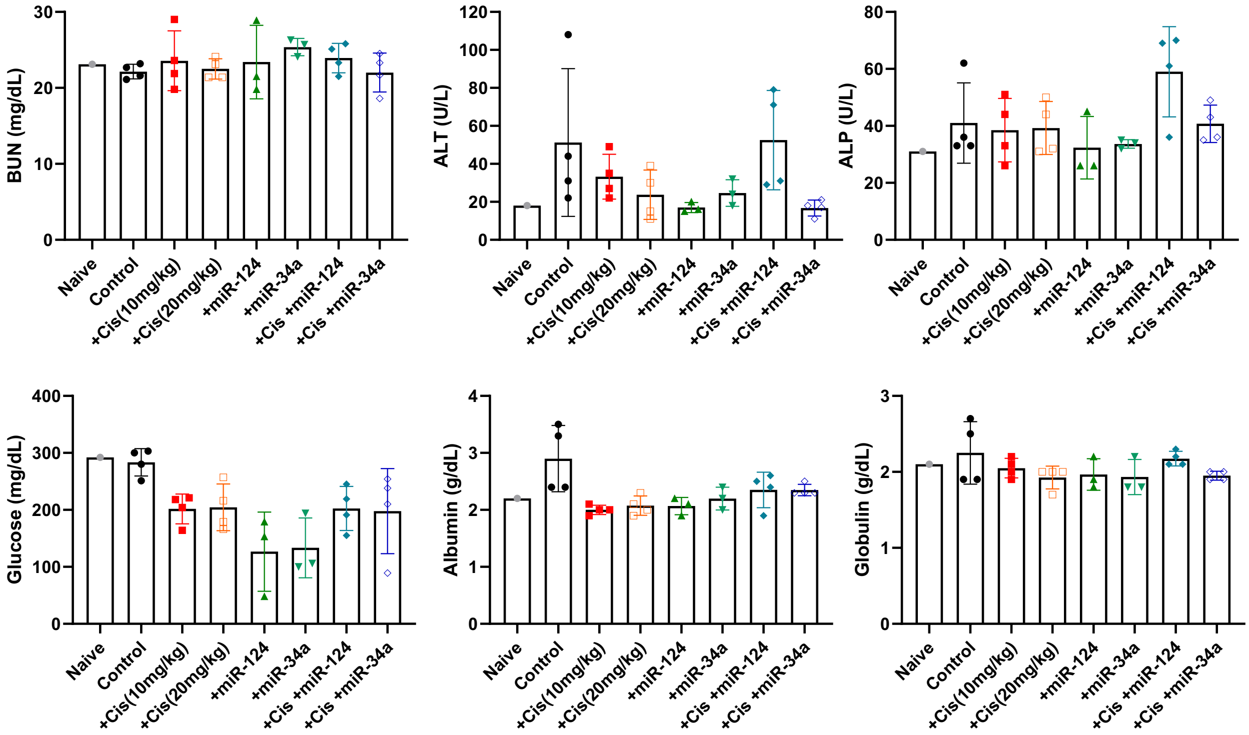


**Supplementary Figure 5. Systemic toxicity assessment of bioRNA and cisplatin treatment in SK-N-SH xenografts.**
(A) Serum chemistry analysis of blood collected at study termination. Renal function (BUN), liver injury markers (ALT and ALP), metabolic parameters (glucose), and serum protein levels (albumin and globulin) were measured to evaluate treatment-associated toxicity. Data are presented as mean ± SD.

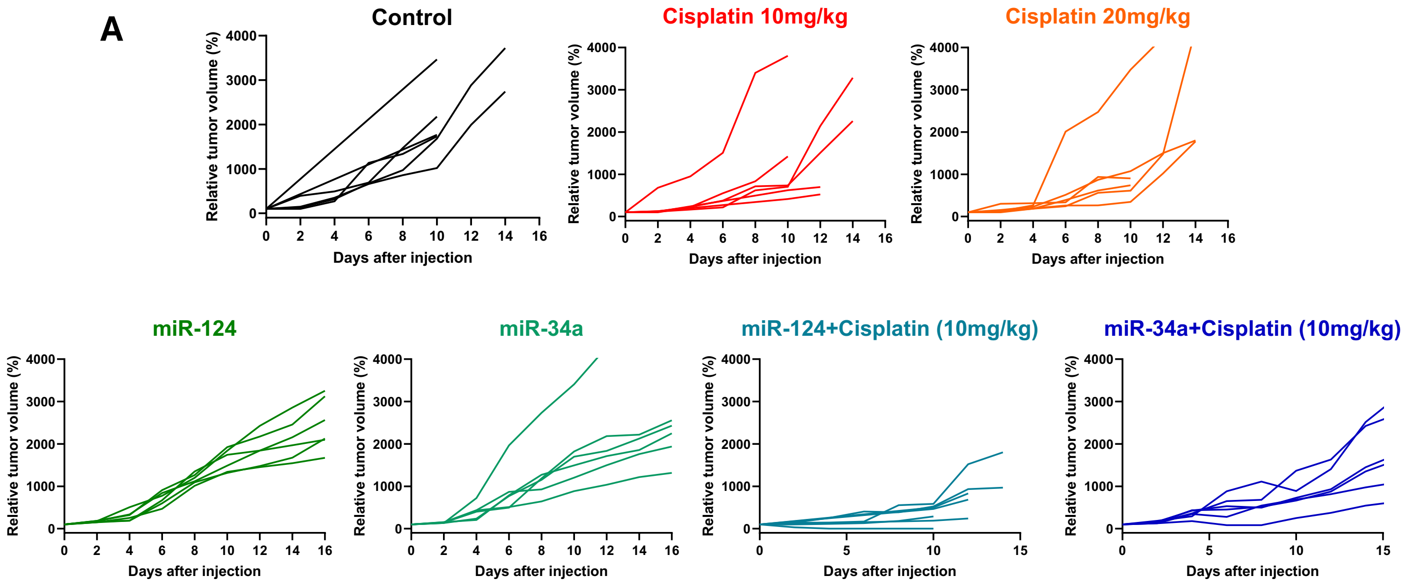


**Supplementary Figure 6. Individual tumor growth trajectories in Be(2)-C xenografts.**
(A) Spider plots showing relative tumor volume progression for individual animals receiving control, cisplatin, bioRNA monotherapy, or combination treatment.

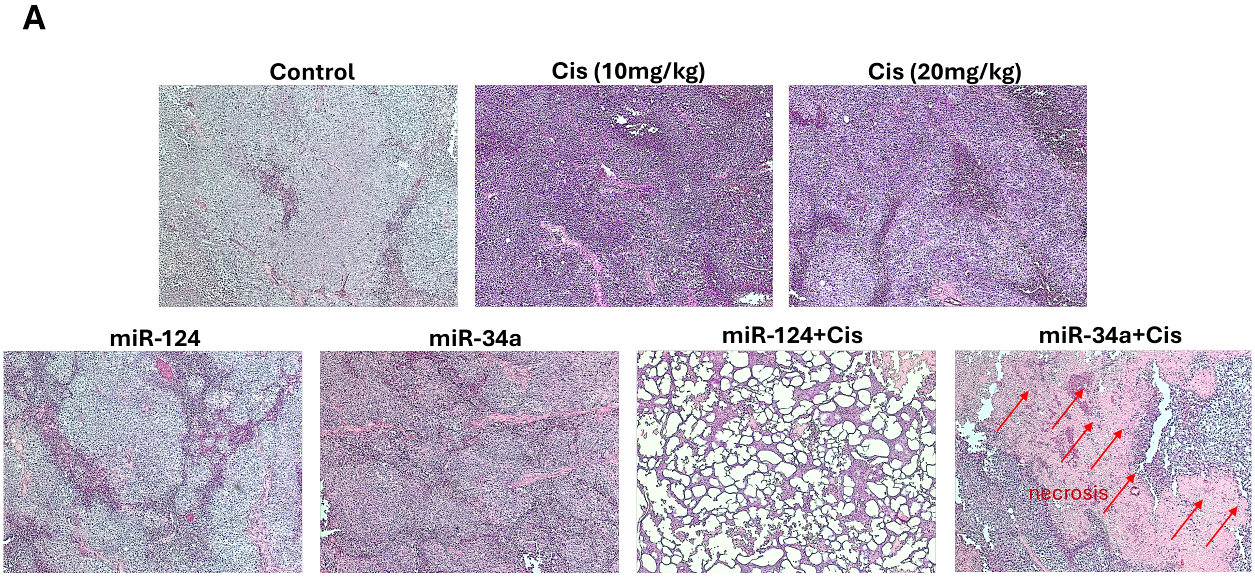


**Supplementary Figure 7. Histopathological evaluation of treatment responses in Be(2)-C xenografts.**
(A) Representative H&E-stained tumor sections collected at study endpoint. Arrows indicate regions of tissue degeneration and necrosis. Combination-treated tumors exhibited extensive tissue destruction and necrotic remodeling.

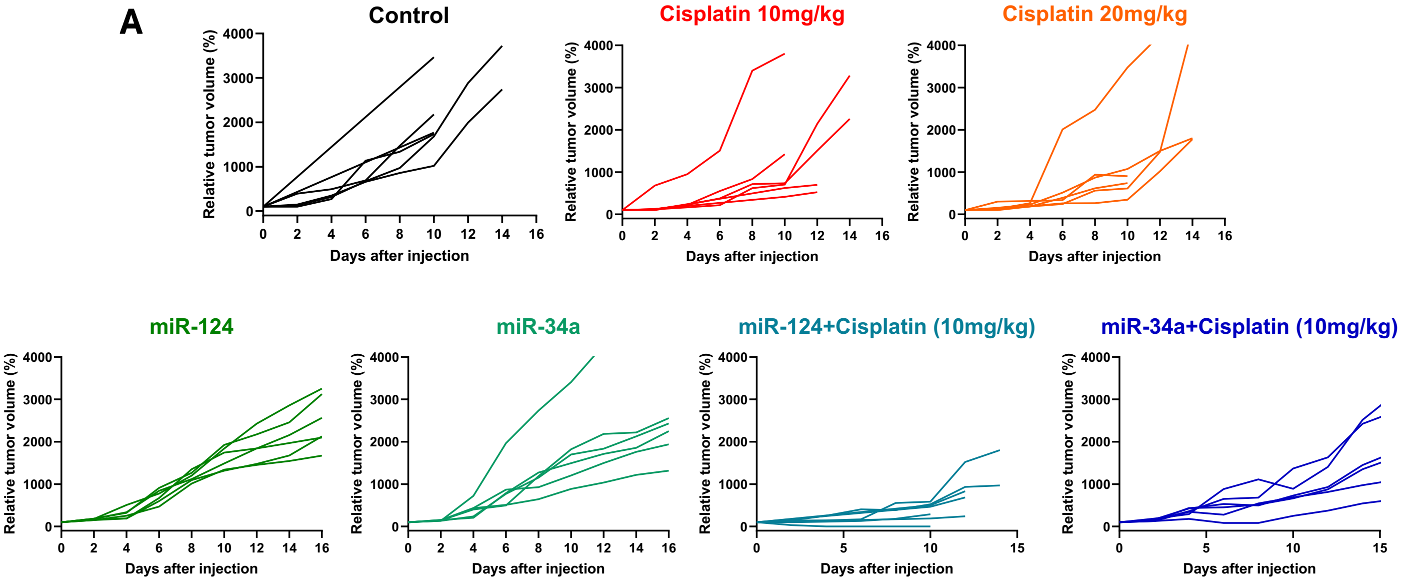


**Supplementary Figure 8. Individual tumor growth trajectories in NB1643 xenografts.**
(A) Spider plots showing relative tumor volume progression for individual animals receiving control, cisplatin, bioRNA monotherapy, or combination treatment.
